# *In vivo* quantification of anterior and posterior chamber volumes in mice: implications for aqueous humor dynamics

**DOI:** 10.1101/2024.07.24.604989

**Authors:** Daniel Kim, Raymond Fang, Pengpeng Zhang, Cheng Sun, Guorong Li, Christa Montgomery, Simon W.M. John, W. Daniel Stamer, Hao F. Zhang, C. Ross Ethier

## Abstract

**Purpose:** Aqueous humor inflow rate, a key parameter influencing aqueous humor dynamics, is typically measured by fluorophotometery. Analyzing fluorophotometric data depends, *inter alia*, on the volume of aqueous humor in the anterior, but not the posterior, chamber. Previous fluorophotometric studies of aqueous inflow rate in mice have assumed the ratio of anterior:posterior volumes in mice to be similar to those in humans. Our goal was to measure anterior and posterior chamber volumes in mice to facilitate better estimates of aqueous inflow rates.

**Methods:** We used standard near-infrared optical coherence tomography (OCT) and robotic visible-light OCT (vis-OCT) to visualize, reconstruct and quantify the volumes of the anterior and posterior chambers of the mouse eye *in vivo*. We used histology and micro-CT scans to validate relevant landmarks from *ex vivo* tissues to facilitate *in vivo* measurement.

**Results:** Posterior chamber volume is 1.1 times the anterior chamber volume in BALB/cAnNCrl mice, i.e. the anterior chamber constitutes about 47% of the total aqueous humor volume, which is very dissimilar to the situation in humans. Anterior chamber volumes in 2-month-old BALB/cAnNCrl and 7-month-old C57BL6/J mice were 1.55 ± 0.36 µL (n=10) and 2.41 ± 0.29 µL (n=8), respectively. This implies that previous studies likely over-estimated aqueous inflow rate by approximately two-fold.

**Conclusions:** It is necessary to reassess previously reported estimates of aqueous inflow rates, and thus aqueous humor dynamics in the mouse. For example, we now estimate that only 0-15% of aqueous humor drains via the pressure-independent (unconventional) route, similar to that seen in humans and monkeys.

## 1. Introduction

Aqueous humor dynamics (AHD) determine intraocular pressure (IOP) and are thus important in understanding ocular physiology and pathophysiology, as well as drug delivery in the anterior segment ^1–4^. Mice are widely used to study AHD, where they show important similarities to humans. Despite the small size of the mouse eye, high-quality measurements are available for a number of important AHD parameters such as murine IOP and outflow facility ^3, 5–8^. However, a key parameter that has received less attention in the mouse eye is the aqueous production rate. The most recent measurements by Toris et al. ^9^ used a customized fluorophotometer to measure inflow in CD-1 mice; other authors have used similar tracer dilution methods ^8, 10, 11^.

Tracer dilution methods, including fluorophotometery, are fundamentally based on measuring the rate of loss of tracer signal in the cornea and anterior chamber and relating this quantity to aqueous inflow rate. This procedure requires accurate knowledge of the corneal and anterior chamber volumes. In the human eye, nearly 80% of total aqueous humor volume is in the anterior chamber ^12^, which has also been assumed to be the case in the mouse eye. However, considering the anatomical differences between the mouse and human eye (e.g. the murine eye has a relatively much larger lens), this assumption may be incorrect. For example, tracer dilution studies in the mouse eye have assumed that anterior chamber volume is approximately equal to <u>total</u> aqueous volume determined by aspiration of all aqueous humor (5.1-7.2 uL) ^10, 11, 13^. If erroneous, this estimate leads to significant errors in determining mouse aqueous inflow rate. Incomplete knowledge of chamber volumes has other implications; for example, when conducting preclinical studies of agents delivered into the mouse eye intracamerally ^14–16^, dosing and dilution effects can be mis-estimated.

Due to the small size of the mouse eye, quantifying anterior and posterior chamber volumes is not trivial. For example, *in vivo* imaging methods such as MRI and ultrasound have insufficient resolution in mice. *Ex vivo* studies of chamber volume are not ideal since tissue handling can deform the globe and lead to incorrect estimates of volumes. To overcome these shortcomings, we adopted an approach based on optical coherence tomography (OCT), informed by *post mortem* studies. OCT has been used to measure anterior chamber depth but not to reconstruct the 3D anatomy of the anterior and posterior chambers ^17–23^. Further, clear boundaries for the posterior chamber cannot be well resolved within a single optical field of view with the OCT beam orientated along the optical axis of the eye, making posterior chamber volume measurement even more challenging. An ideal solution is to obtain a volumetric OCT image of the entire anterior and posterior chambers *in vivo* and measure chamber volumes from the reconstructed volumetric image. Towards this end we used a robotic visible-light OCT (vis-OCT) system. Vis-OCT has a higher axial resolution than conventional OCT using near-infrared light, with an axial resolution ∼ 1.3 microns in tissue, ^24^ allowing us to reconstruct the anterior segment with high resolution *in vivo*. We validated the accuracy of volume measurements and overall volumetric reconstruction with a 3D-printed phantom, and then segmented robotic vis-OCT images, with landmarks validated by other imaging modalities, to obtain chamber volumes in living mice.

## 2. Methods and Materials

### 2.1 Animal handling

All procedures were approved by the relevant institutions’ Institutional Animal Care and Use Committee and conformed to the Association for Research in Vision and Ophthalmology Statement on Animal Research. All mice used in this study were wild type.

At Northwestern University, vis-OCT imaging was carried out in ten 2 month-old adult BALB/cAnNCrl albino mice (Charles River Laboratories, Skokie, IL; 5 males, 5 females). Using albino mice eliminated pigment-induced scattering, providing the best possible visualization of structures posterior to the iris. Body weights of BALB/cAnNCrl mice were not measured. For strain comparison, we also imaged the anterior chambers of eight 7-month-old C57BL/6J mice (JAX stock number 000664, Jackson Laboratory, Bar Harbor, Maine; 4 males, 4 females). Mean body weights for C57BL/6J mice were 35.5g (males) and 28.0g (females). All mice were housed under 12h:12h light:dark cycles within the Center for Comparative Medicine at Northwestern University.

At Columbia University, C57BL/6J mice (JAX stock number 000664) were obtained from Jackson Laboratory (Bar Harbor, Maine) while DBA/2J mice were wild type mice from a DBA/2JSj substrain that we had separated from The Jackson laboratory’s DBA/2J strain in 2019 (and so are essentially the same genetically). Animals were housed under a 14h:10h alternating light:dark cycle within the Institute of Comparative Medicine animal facility.

At Duke University, mice were handled in accordance with approved protocol (A226-21-11). C57BL/6J mice were purchased from the Jackson Laboratory (JAX stock number 000664) and CD-1 mice were purchased from Charles River (Charleston, SC; stock number 22), bred/housed in clear cages and kept in housing rooms at 21°C on a 12h:12h light: dark cycle in the Duke animal facility.

### 2.2 Histology and conventional near-infrared OCT imaging

Initial studies were conducted at Duke University. C57BL/6J mice were anesthetized using isoflurane. Once they reached a deep plane of anesthesia, animals were decapitated, and eyes were carefully enucleated and immersion fixed in 4% PFA + 1% glutaraldehyde at 4 °C. The posterior sclera and part of the retina were carefully dissected, after which the remainder of the globe was processed for embedding in Epon using standard approaches. The block was then trimmed, oriented and sectioned until the sectioning plane reached the approximate center of the eye. Sections were then collected, stained with 1% methylene blue and examined by light microscopy (Axioplan2, Carl Zeiss MicroImaging, Thornwood, NY).

To conduct conventional OCT imaging, mice were anesthetized with ketamine (100 mg/kg)/xylazine (10 mg/kg) and secured in a custom-made platform. Eyes were imaged with an Envisu R2200 high-resolution spectral domain (SD)-OCT system (Bioptigen Inc., Research Triangle Park, NC). The mouse was positioned until the OCT probe faced the center of the cornea, and cross-sectional images spanning from the nasal to temple sides of the globe were recorded.

### 2.3 Micro-CT image acquisition

All micro-CT imaging experiments were conducted at Columbia University. Eyes were collected within 15 minutes of euthanasia by cervical dislocation and immersion fixed overnight in 3% paraformaldehyde plus 1% glutaraldehyde in phosphate-buffered saline at 4 °C. Whole eyes were stained either with eosin-Y as previously described ^25^ and phosphotungstic acid (PTA), or with PTA alone as described previously ^26^ then dehydrated with hexamethyldisilazane. Eyes were scanned in a Bruker SkyScan 2214 multiscale-CT system (Micro Photonics Inc., Allentown, PA) utilizing a tungsten x-ray source at 53 keV and an 11 Mpixel CCD detector. This setup has an achievable voxel size of 120 nm and maximum spatial resolution of 500 nm. A 360-degree scan was acquired with rotation steps of 0.15 degrees and 6 frame averages. Projection images were reconstructed with Bruker NRecon software. Three-dimensional virtual sections for the figures presented were produced with Bruker’s CTvox software. Opacity and luminosity were adjusted for each image to show the target structures as appropriate.

### 2.4 Vis-OCT image acquisition

All vis-OCT imaging experiments were conducted at Northwestern University. Before imaging, mice were anesthetized with a ketamine/xylazine cocktail (ketamine: 11.45 mg/mL; xylazine: 1.7mg/mL, in saline) delivered via intraperitoneal injection (10 mL/kg). During imaging, we maintained the mouse’s body temperature using a heating lamp and applied artificial tears to prevent corneal dehydration.

To obtain high-resolution *in vivo* anterior segment images, we used an experimental robotic vis-OCT system ^27^, technical details of which are given in the supplemental material (Figure 1). Due to the location of the posterior chamber within the anterior segment and the angle-dependent backscattering of the lens and borders of the posterior chamber, the boundaries of the posterior chamber are best visualized when the incident OCT beam is normal to the limbus. However, in this configuration, OCT does not have sufficient imaging depth to capture the entire anterior segment within a single volumetric acquisition. To reconstruct the entire anterior segment, we captured eight volumes (Fig. 1c), each separated by 45 degrees, around the eye positioned by a 6-degree-of-freedom robotic arm (Mecademic, Montreal, Canada; Meca500 Robot Arm). Each scan consisted of 512 B-scans, and each B-scan consisted of 512 A-lines, together acquiring a volume with lateral area 2.04 × 2.04 mm^2^ and depth 1.56 mm in air. Adjacent volumetric scans had an overlap of approximately 40%. We acquired each volume using a temporal speckle averaging scan pattern, where each B-scan was repeated twice per volume, and we acquired two repeated volumes at each of the eight positions ^28^. We also processed the interferograms using the optical microangiography algorithm to generate visible-light OCT angiography (vis-OCTA) images ^29^.

**Fig. 1.**
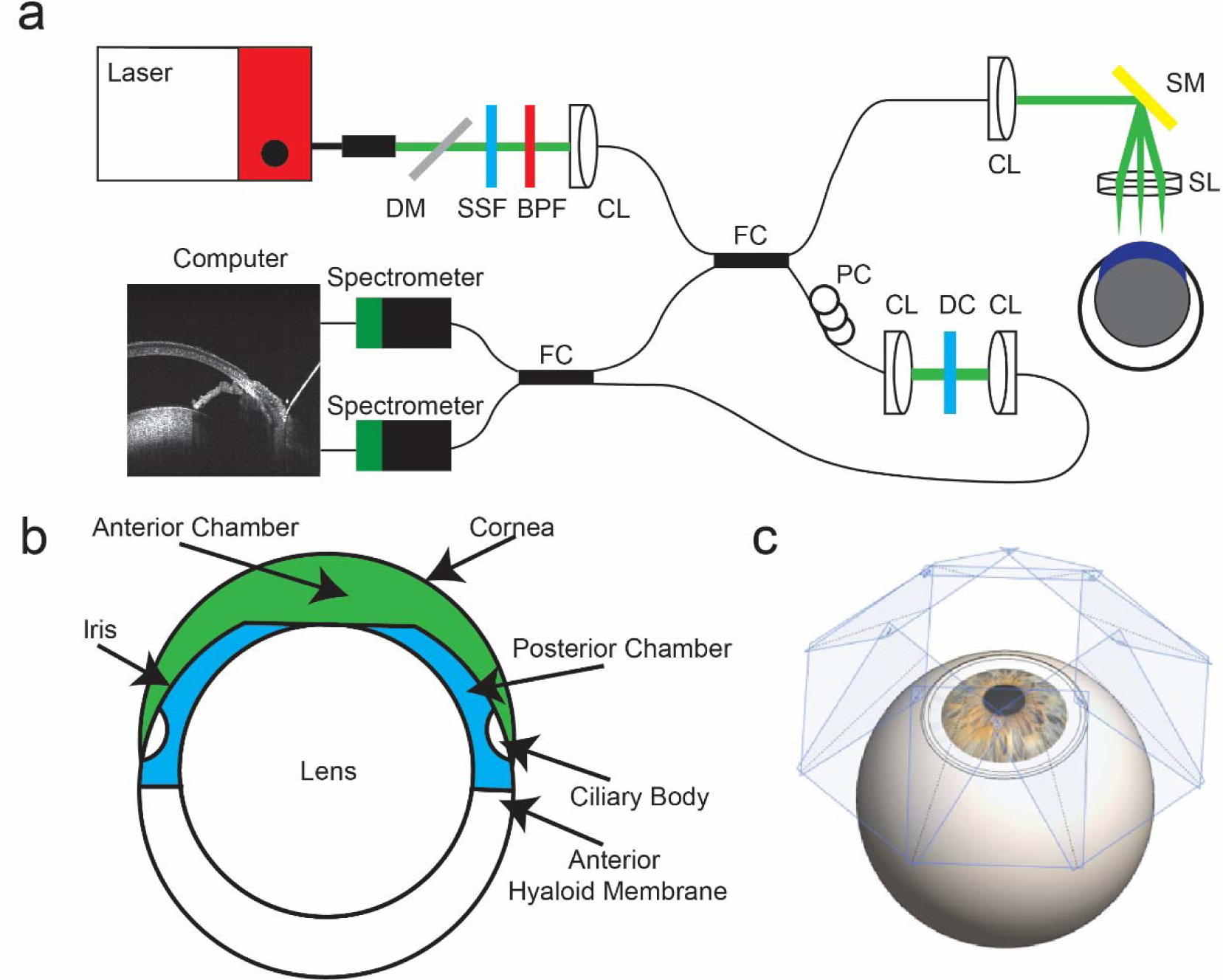
Experimental setup of vis-OCT imaging. **a)** Schematic of robotic vis-OCT. Light from an NKT laser is filtered by a dichroic mirror (DM), spectral shaping filter (SSF), and bandpass filter (BPF). The output light is coupled into a collimator (CL) and split by a 90:10 fiber coupler (FC). The reference arm includes a polarization controller (PC) and dispersion compensation (DC). Light in the sample arm is scanned by a galvanometer scanning mirror (SM) before being focused by a 25-mm scan lens (SL). The interference signal is split by a 50:50 fiber coupler (FC) into two spectrometers. **b)** Schematic cross-section of the mouse eye with the anterior chamber shaded in green and the posterior chamber in blue. **c)** Eight vis-OCT volumes, with scan planes perpendicular to the incident vis-OCT beam shaded in blue, are acquired around the eye.

### 2.5 Vis-OCT volume fusion

To combine the eight volumes, we developed an algorithm based on point cloud registration methods commonly used in LIDAR ^30, 31^. Briefly, we represented each volume as a point cloud and registered the point clouds together. When two sets of point clouds had sufficient overlap, we found the transformation matrix that minimized the distance between the overlapping point clouds ^27^ (Fig. 2).

**Fig. 2.**
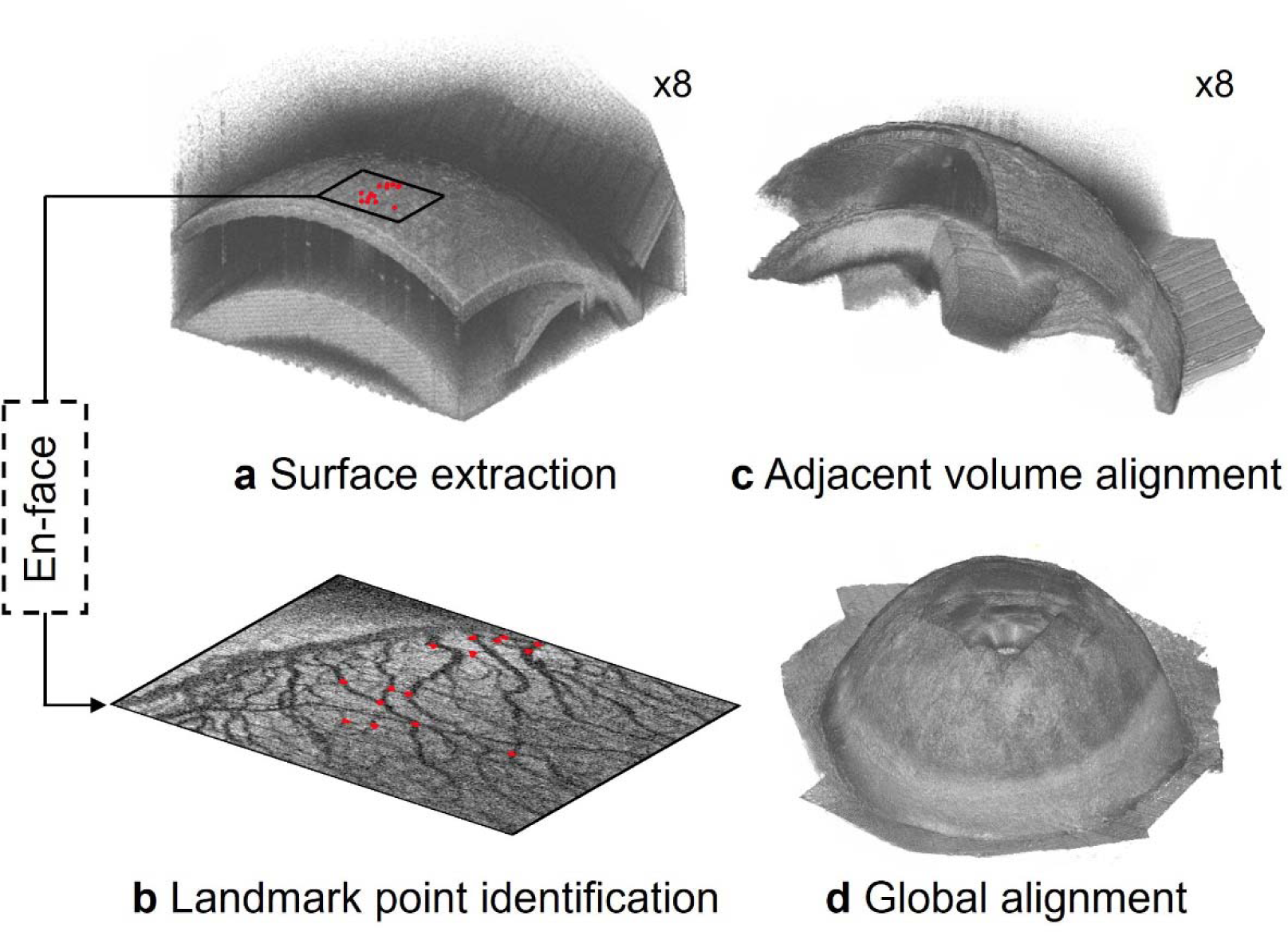
Overview of anterior segment reconstruction. **a)** The outer surface of the globe for each sub-volume is extracted and **b)** landmark points are identified along the top surface of each sub-volume. The locations of several landmark points are shown by the red dots, whose spatial locations correspond to the red dots in a). **c)** The landmark points are registered between adjacent volumes and used to align them. **d)** After alignment, all sub-volumes are mapped into a common spatial reference frame.

To transform each sub-volume into a common spatial reference frame, we represented the outer surface of the eye for each volume as a point cloud. To determine the spatial position of the outer surface, we applied a threshold to each vis-OCT volume, kept the largest connected binarized object, and found the outer surface of the remaining binarized object (Fig. 2a). For vis-OCT volumes acquired adjacent to each other, we identified common reference points (landmarks) in the overlapping regions of the point clouds. Specifically, from vis-OCTA images, we manually identified 15 blood vessel branch points as landmarks (Fig. 2b). We then used the M-estimator sample consensus (MSAC) algorithm to obtain an initial estimate of the rigid transformation matrix that would match the spatial position of the landmark points between two adjacent volumes in three dimensions ^32^ (Fig. 2c). We aligned all volumes within a common coordinate system and obtained the pixelwise intensity of that combined volume (Fig. 2d), as described in the supplemental material.

### 2.6 Anterior and posterior chamber segmentation and volumetric rendering

We defined the anterior chamber as the space bounded by the cornea, anterior iris, and anterior lens and the posterior chamber as the space between the lens, posterior iris, and anterior hyaloid membrane ^33^ (Figure 1b). To generate volumetric representations of the anterior and posterior chambers, we used the Segment Anything Model (SAM; Meta AI, New York City, NY ^34^) to segment individual B-scan images and combine the segmented results to form a volumetric representation of the chambers. SAM is a general segmentation model that allows zero-shot segmentation of various images ^34^. With appropriate fine-tuning steps, SAM has previously been shown to be compatible with medical images, including applications in CT, MRI, and OCT ^35^.

To segment the anterior and posterior chambers, we first obtained an initial mask of the chambers using SAM. During this step, we manually marked points every 50 B-scans within and outside the chambers as SAM’s zero-shot segmentation requires users to identify inlier and outlier regions with point inputs. We interpolated the position of the inlier and outlier points with a third-order polynomial to approximate their locations across the B-scans and fed these inputs to the model checkpoint based on the Vision Transformer-Huge (ViT-H) image encoder ^36^. Following initial segmentation, we manually checked and fine-tuned the segmentations using the AI segmentation website Biodock ^37^. With fine-tuning, the output neural network generated high-quality segmentations of the chambers without user input.

Fig. 3 illustrates the process of reconstructing the posterior and anterior chambers. Each vis-OCT volume consists of 512 B-scans, from which we segmented the posterior chamber (blue-shaded region in Fig. 3a) for each B-scan. Then, we merged the segmented posterior chamber for each of the eight vis-OCT volumes (Fig. 3b). Finally, we applied the rigid transformation matrices obtained during montaging to map the posterior chamber for each vis-OCT volume into a common reference frame to reconstruct the entire volume as shown in Fig. 3c. To reconstruct the anterior chamber, we used the fully reconstructed montaged anterior segment volume generated using the methodology described in section 2.5 (Fig. 3d). Next, we used the trained SAM to segment the anterior chamber in each digital cross-sectional image of the montaged volume along the x-z plane, as highlighted by the green shaded region in Fig. 3e. Finally, we merged the segmented anterior chamber in each digital cross-sectional image to form the circumlimbal volume of the anterior chamber (Fig. 3f). We measured the volumes of each chamber in each eye after segmentation based on the vis-OCT voxel size and the number of voxels. We determined the voxel size in the axial direction based on the parameters of the spectrometers^38^ and the lateral direction by calibrating to a gridded sample with known grid sizes (R1L3S3P, Thorlabs, Newton, NJ).

**Fig. 3.**
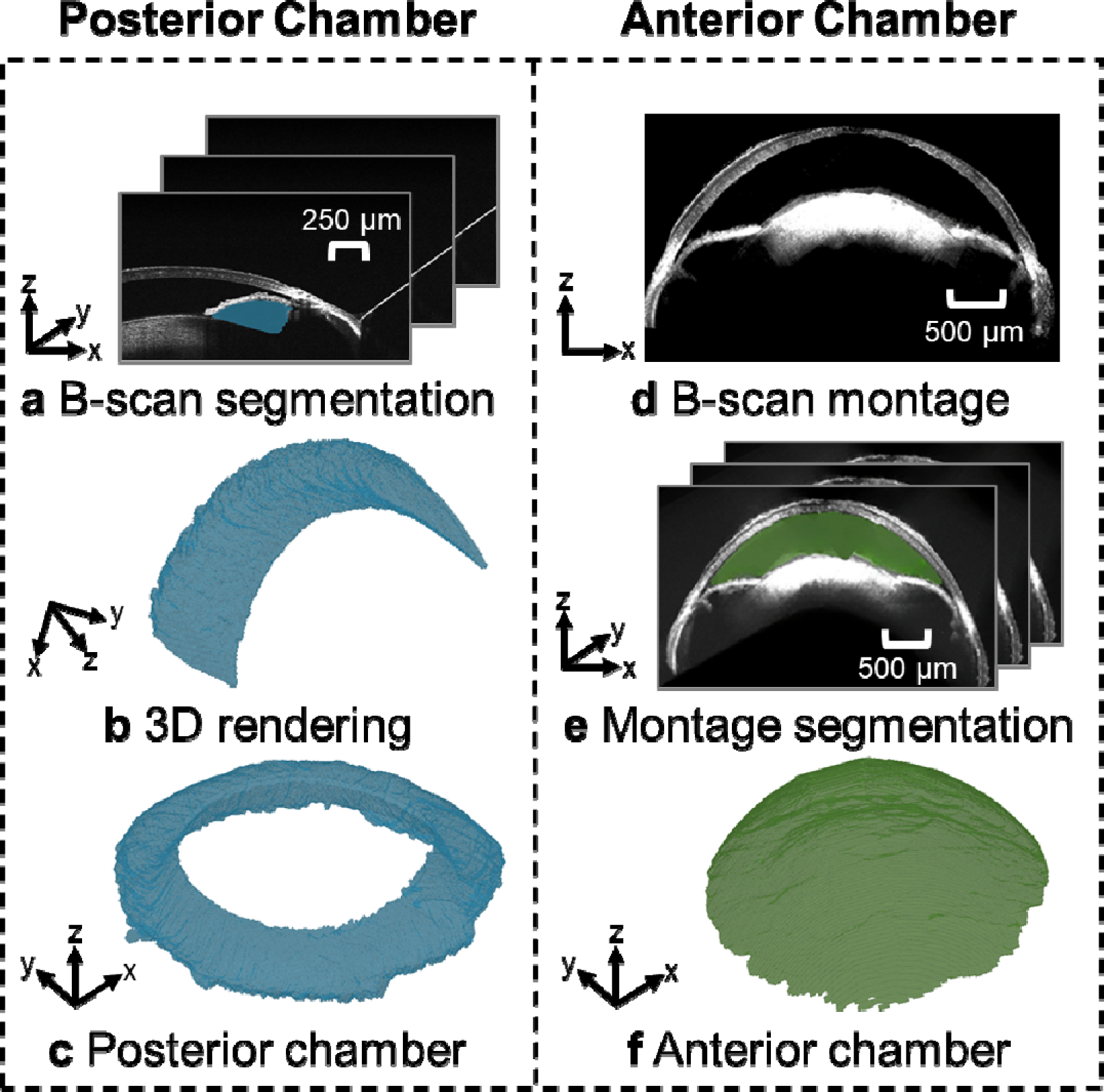
Posterior chamber and anterior chamber reconstruction processes. **a)** B-scans were segmented using SAM, with the posterior chamber shaded in blue. **b)** After segmenting all B-scans, we generated a volumetric representation of the posterior chamber for each volume. A transformation matrix mapped each volume into a common coordinate system and the union of **c)** segmented volumes was taken to be the posterior chamber. **d)** The transformation matrices were also used to map the OCT structural data into a common reference frame, where **e)** the anterior segment of each cross-section was segmented using SAM. **f)** The union of the segmented B-scans was used to generate a volumetric representation of the anterior chamber.

### 2.7 Posterior chamber volume correction

Due to its position within the eye and its tenuous structure, the anterior hyaloid membrane – needed to define the posterior boundary of the posterior chamber – could not reliably be visualized by vis-OCT, even in BALB/cAnNCrl mice. Thus, the posterior boundary of the posterior chamber segmentation generated by SAM was a curve connecting the lens with the ciliary body. However, the ciliary body lies anterior to the hyaloid membrane (Supplementary Figure S2), so the anterior hyaloid membrane location generated by SAM was incorrect and led to an underestimation of posterior chamber volume. To correct this underestimation, we approximated the posterior boundary of the posterior chamber (anterior hyaloid membrane) by the equator of the eye ^39^, based on our micro-CT images. As described in detail in the Supplemental Materials, we thus approximated the anterior hyaloid membrane location by the plane that passed through the center of the lens and that was normal to the optical axis of the eye (Fig. 4).

**Fig. 4.**
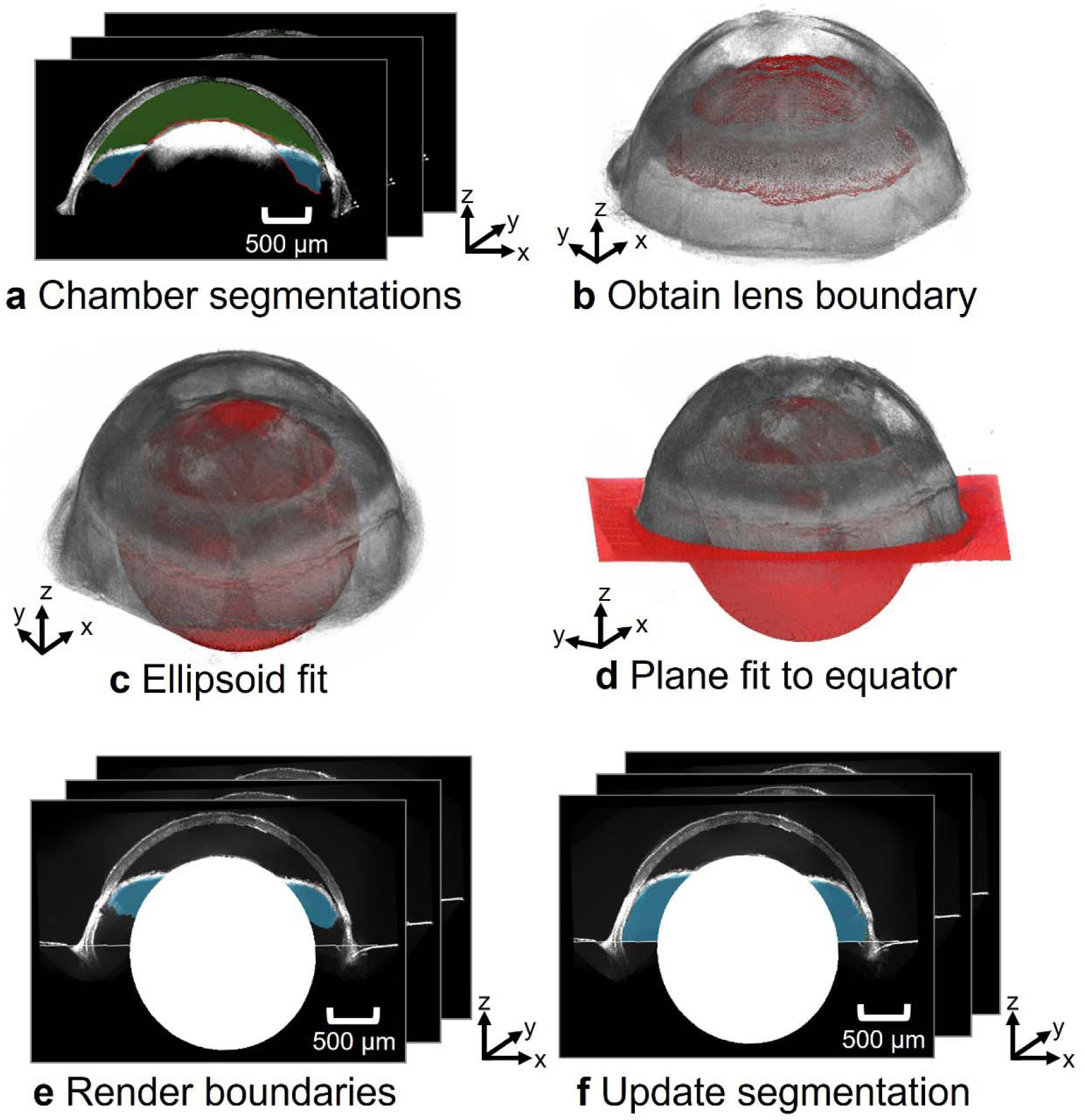
Workflow for identifying the posterior boundary of posterior chamber. **a)** Chamber segmentations were used to obtain the posterior boundary of the anterior chamber (green) and the interior boundary of the posterior chamber (blue). **b)** These boundaries were taken to coincide with the anterior surface of the lens. **c)** An ellipsoid (red) was fit to the lens’s upper boundaries. The center of the ellipsoid and the optical axis of the eye were used to **d)** generate a plane at the equator of the eye, approximating the location of the anterior hyaloid membrane. **e)** The posterior border of the reconstructed posterior chamber in the montaged B-scans is **f)** updated with the estimated position of the anterior hyaloid membrane.

### 2.8 Volume measurement validation

We validated our vis-OCT volume measurement using a 3D-printed phantom consisting of a hemisphere with a cavity to mimic the posterior chamber of the mouse eye (Fig. 5a). We designed the phantom using SolidWorks^TM^ and 3D-printed the phantom using our homemade micro continuous liquid interface production (µCLIP) system ^40^ with photocurable clear resin, which was made by mixing 98.85 wt.% Poly(ethylene glycol) diacrylate (PEGDA, Sigma-Aldrich Inc., Saint Louis, MO) as a low-viscosity monomer, 1 wt.% Phenylbis (2,4,6-trimethylbenzoyl), phosphine oxide (Irgacure 819, Sigma-Aldrich Inc., Saint Louis, MO) as photoinitiator, and 0.15 wt.% Avobenzone (Tokyo Chemical Industry Co., Tokyo, Japan) as UV absorber. After printing, we washed the phantom with isopropyl alcohol to remove any remaining resin and post-cured it under UV light. Finally, we filled positive contrast resin into the 3D-printed hollow phantom structure and placed the phantom under UV light to fully cure the contrast resin. The positive contrast resin consisted of 93.8 wt.% 2-Hydroxyethyl methacrylate (HEMA, Sigma-Aldrich Inc.,Saint Louis, MO), 3 wt.% Ethylene glycol dimethacrylate (EGDEA, Sigma-Aldrich Inc., Saint Louis, MO), 2.2 wt.% Irgacure 819, and 1% Intralipid (Sigma Aldrich, Saint Louis, MO). We visualized the geometry of the 3D-printed phantom by scanning electron microscopy (SEM) and found the volume of the hollow region by filling this region with de-ionized water and measuring the mass of the phantom before and after water filling. To conduct SEM imaging, we deposited a thin layer of 10-nm Au/Pd onto the printed sample with sputter coating (Denton Vacuum, Moorestown, NJ) and acquired images using an EPIC SEM FEI Quanta 650 (FEI, Hillsboro, OR). We measured the phantom volume from eight vis-OCT volumes using the methodology described in section 2.6.

**Fig. 5.**
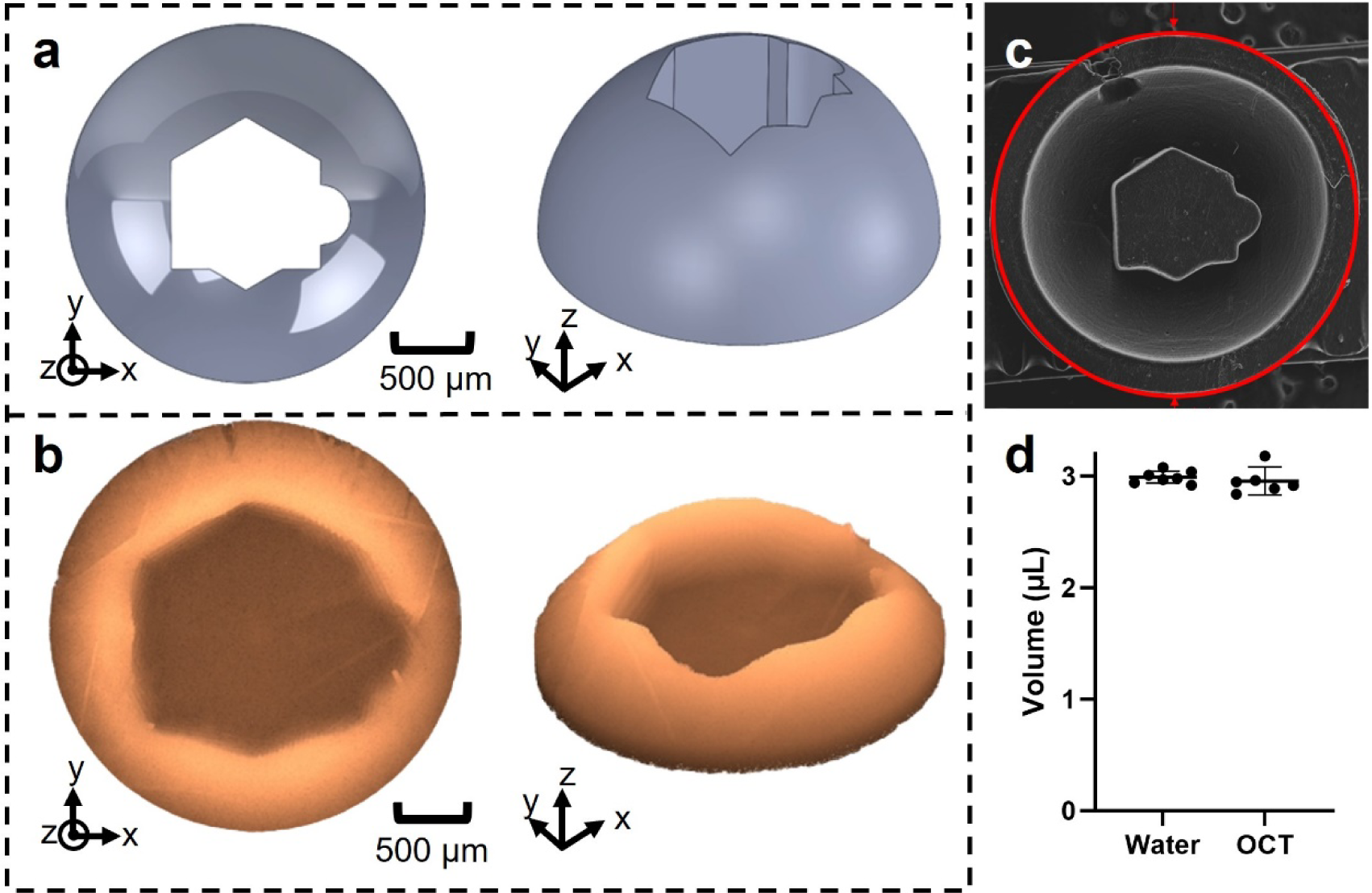
Validation of reconstruction volume accuracy. **a)** Design of phantom cavity within CAD. **b)** Reconstructed phantom volume using vis-OCT after montaging. **c)** SEM image of the 3D-printed phantom. **d)** The volumes of the phantom obtained by water weighing and from the OCT reconstruction agreed to within one percent, with error bars representing the 95% confidence intervals. (*P < 0.05; **P< 0.01, ***P < 0.001, ****P < 0.0001).

### 2.9 Statistical analysis

All statistical computations were carried out using GraphPad Prism 10.1.0 (Boston MA). We compared the phantom volumes measured using water weight vs. OCT with an unpaired t-test. Similarly, we compared anterior and posterior chamber volumes as well as anterior chamber volume between BALB/cAnNCrl and C57BL/6J mice using an unpaired t-test. All numerical data are presented as mean ± SD. The error bars on all plots represent 95% confidence intervals. We used p < 0.05 as the threshold for statistical significance.

## 3. Results

### 3.1 Histology, conventional OCT imaging and micro-CT imaging

Although significant distortion was evident is histologic images (Figure S1), a morphometric analysis based on measuring anterior and posterior chamber boundaries and rotating the images through 180 degrees to compute the corresponding volumes suggested that anterior and posterior chamber volumes were approximately equal. To investigate the situation *in vivo*, we thus carried out conventional near-infrared (NIR) OCT imaging and a similar morphometric analysis, rotating the OCT images through 180 degrees and estimating anterior and posterior chamber volumes, again finding that < 50% of total aqueous humor volume resided within the anterior chamber (data not shown).

The above results were suggestive but not definitive due to tissue deformation occurring during histologic processing and poor visualization of the posterior chamber structures by conventional OCT imaging. We therefore undertook micro-computed tomographic (micro-CT) imaging of *post mortem* eyes designed to more clearly identify the location of posterior chamber structures, particularly the anterior hyaloid membrane. We observed (Supplemental Figure S2) that the anterior hyaloid membrane, which we took as the posterior margin of the posterior chamber, was approximately located at the equator of the eye. Importantly, we observed that the position of the anterior hyaloid membrane was similar in both 13-month-old DBA/2J and 1.5-month-old C57BL/6J mice (Supplementary Fig. S2), suggesting that the equator was an appropriate landmark for the anterior hyaloid membrane.

### 3.2 Algorithm validation using a phantom

Before using very high spatial resolution vis-OCT imaging to quantify the anterior and posterior chamber volumes *in vivo*, we assessed the accuracy of our vis-OCT volume measurement and reconstruction algorithm by imaging a 3D-printed phantom containing a cavity mimicking the posterior chamber (Fig. 5b). The outer edge of the printed phantom had a diameter of 3 mm, approximately the diameter of the mouse eye. Since 3D printing is subject to errors when printing features with sub-millimeter dimensions, we acquired an SEM image of the phantom to validate its structure and dimensions (Fig. 5c). We found that the features of the SEM image matched those of our reconstructed vis-OCT image. We then measured the cavity volume by determining the mass of water required to fill the phantom cavity, obtaining 2.99 ± 0.06 µL (n=7 technical replicates). The volume determined by vis-OCT imaging was 2.96 ± 0.12 µL (n=6 technical replicates), which was within 1% of the volume determined by the water filling approach (Fig. 5d). This difference was not statistically significant, and we conclude that our vis-OCT-based approach accurately determined the volume of a cavity in a phantom of similar size to the anterior and posterior chambers.

### 3.3 In vivo anterior and posterior chamber volume measurements

We reconstructed the entire anterior and posterior chambers of 2-month-old BALB/cAnNCrl albino mice (n=10; Fig. 6a). As expected, the anterior chamber formed a continuous volume anterior to the iris, while the posterior chamber formed a continuous volume posterior to the iris. The anterior chamber resembled a spherical cap below the cornea, and the posterior chamber resembled the upper half of a torus. When viewing the volumetric reconstruction from the posterior view (Fig. 6b), we found that the outer radius of the posterior chamber was larger than the anterior chamber. Fig. 6c shows a cross-sectional view of both chambers.

**Fig. 6.**
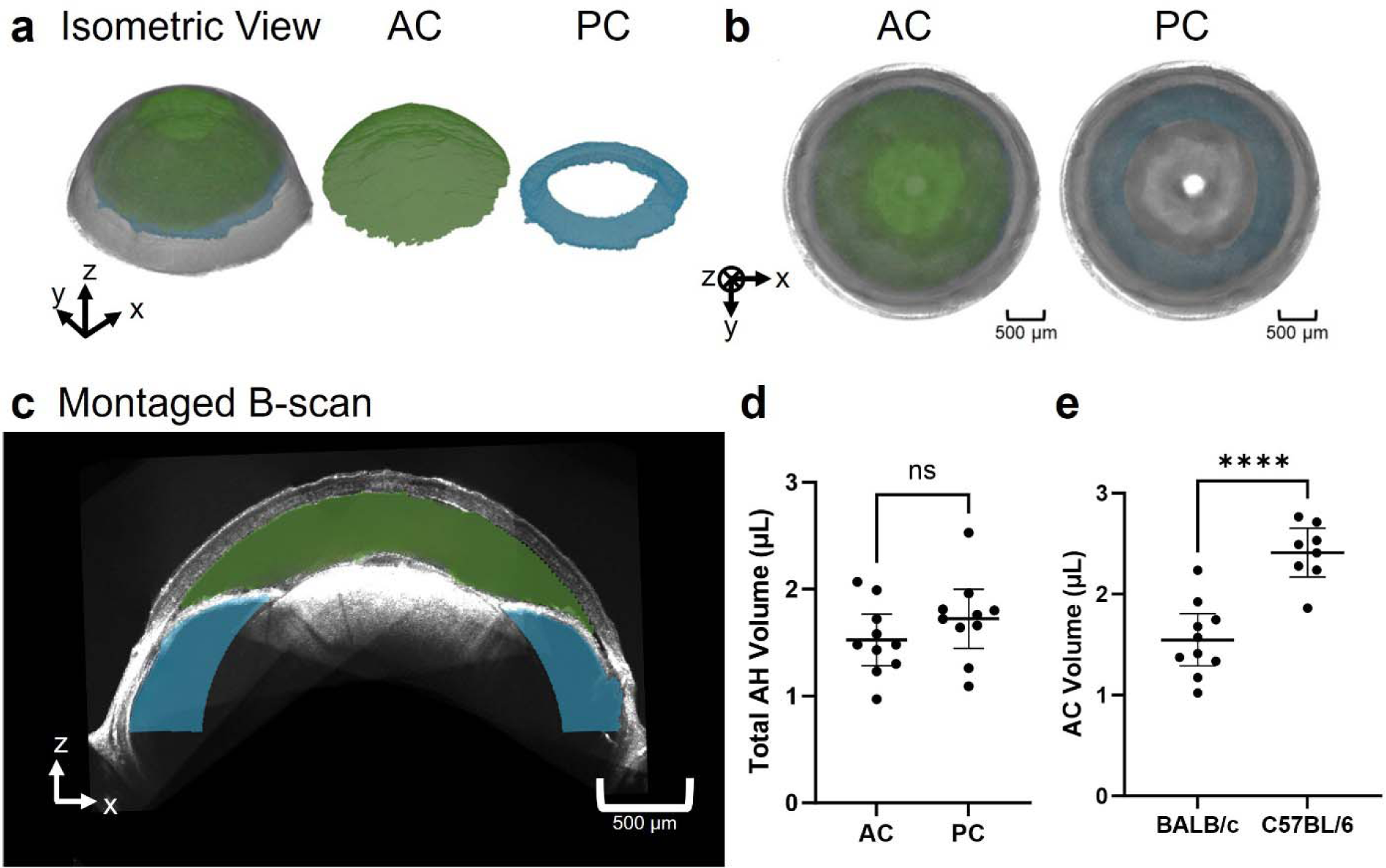
Reconstruction of the anterior and posterior chambers. **a)** Isometric view of the reconstructed anterior segment with anterior chamber (AC) in green and posterior chamber (PC) in blue. **b)** Posterior view of reconstruction with the AC and PC shaded in green and blue respectively. **c)** Montaged B-scan of the anterior segment with the anterior chamber overlayed in green and the posterior chamber in blue. **d)** Comparison of the anterior chamber volume and posterior chamber volume in BALB/cAnNCrl mice reveals that the volumes of aqueous humor in the anterior and posterior chambers are comparable. **e)** Comparison of anterior chamber volumes in 2-month-old BALB/cAnNCrl albino mice and 7-month-old C57BL/6J mice reveals that the anterior chamber is larger in the older C57BL/6J mice. (*P < 0.05; **P< 0.01, ***P < 0.001, ****P < 0.0001).

In 2-month-old BALB/cAnNCrl mice, the measured anterior chamber volume was 1.53 ± 0.34 µL and the posterior chamber volume was 1.72 ± 0.39 µL (Fig. 6d). The total (anterior plus posterior chamber) volume was 3.25 ± 0.49 µL. As an approximate indicator of overall eye size, we also measured the distance between the apexes of the iridocorneal angle on opposite sides of the eye to be 2.87 ± 0.14 mm.

We found that the posterior chamber had a greater volume than the anterior chamber in seven of the ten mouse eyes. Overall, the ratio of the anterior chamber volume to the posterior chamber volume ranged from 0.50 to 1.36, with an average of 0.93 ± 0.28. We found the anterior chamber constitutes 33% to 58% of the total aqueous humor volume (anterior + posterior chamber volumes), with an average of 47 ± 8%.

To confirm that the accuracy of chamber volumes was not impacted by the volumetric montaging algorithm, we measured anterior chamber volume using both the montaged volume consisting of eight volumes and from a single-volume vis-OCT acquisition that captured the entire anterior chamber. Unfortunately, we could not capture the posterior chamber within a single volume, so we only compared the anterior chamber volume measurements. We found an anterior chamber volume of 1.53 ± 0.34 µL from the montaged reconstructions and 1.55 ± 0.36 µL (n=10) from the single-volume acquisitions, with no statistical difference between the two methods. As compared to the single-volume acquisition, the multi-volume reconstructed volumes had greater volume for six eyes and smaller volume for four eyes. We conclude that it is unlikely the reconstruction scheme biased the measured volume in a specific direction.

To investigate chamber volumes in another strain, we also imaged the anterior segments in 7-month-old C57BL/6J mice, measuring an anterior chamber volume of 2.41 ± 0.29 µL (n=8), which was on average 40% larger than the anterior chamber volume measured in 2-month-old BALB/cAnNCrl animals (Fig. 6e). (Note that because the highly pigmented iris of C57BL/6J mice did not allow sufficient light penetration to the posterior chamber, we could not image the posterior chamber in these mice.) Since these animals were older than the BALB/cAnNCrl mice we used, the interpretation of this difference involves both age and strain effects (see Discussion). We found the distance between the apexes of the iridocorneal angle on opposite sides of the eye as 3.27 ± 0.25 mm, which was 14% larger than in the 2-month-old BALB/cAnNCrl mice. If we assume that anterior chamber volume scales with linear dimensions cubed, the 14% difference in linear dimension would correspond to a 48% greater anterior chamber volume in 7-month-old C57BL/6 mice vs. 2-month-old BABL/cAnNCrl mice, which is comparable to, although slightly larger than, the measured 40% difference.

## 4. Discussion and conclusions

In this work, we used robotic vis-OCT imaging to obtain volumetric representations of the anterior and posterior chambers of mice *in vivo*. Two potential applications of this information include a better understanding of aqueous humor dynamics and optimizing intracameral injections in mice. For example, knowledge of inflow rate calculation often depends on anterior chamber volume and is required when using the modified Goldmann’s equation to estimate AHD parameters such as unconventional aqueous drainage rate^41^. Further, several existing models of aqueous humor fluid dynamics, such as the movement of aqueous humor through the iris-lens channel, require knowledge of specific volumes of the chambers ^42^.

A key finding of this work is that only approximately 47% of the total aqueous humor volume is contained within the anterior chamber in BALB/cAnNCrl mice. This is generally consistent with findings based on histology and conventional NIR-OCT, which found that approximately half of the aqueous humor resides in the mouse posterior chamber.

To preliminarily explore the effect of mouse strain, we also imaged the anterior chamber of 7-month-old C57BL/6J mice, obtaining 2.41 ± 0.29 µL compared to 1.55 ± 0.36 µL in the BALB/cAnNCrl animals. At first glance, this might suggest that C57BL/6J mice have larger anterior chambers vs. BALB/cAnNCrl mice; however, we must account for the age difference between the two cohorts. More specifically, globe size is known to increase with age in mice ^43, 44^, with a particularly significant increase over the first 6 months. Using data from Li et al.^43^, we estimate that globe diameter increases by 8% in C57BL/6J mice between 2 and 7 months of age, which implies a 26% increase in globe volume if we assume isotropic growth. This is less than the 40% difference that we observed between 2-month-old BALB/cAnNCrl mice and 7-month-old C57BL/6J mice but suggests that much of the difference in anterior chamber volume we observed was likely an age effect. It is also noteworthy that albinism, present in the BALB/cAnNCrl mice we used in this study, is known to affect IOP and anterior segment development^45^. Thus, further studies will be required to evaluate how age, sex and strain affect anterior chamber volume. Future studies should also consider variables such as type of anesthesia and hydration status.

It is important to compare our measured volumes with previous reports. We are not aware of any papers describing direct measurements of anterior chamber volume in mice, other than a passing comment by Avila et al.^8^, who stated “we have estimated the anterior chamber volume to be approximately 2 _μ_l, calculated as the volume of revolution from the projection of a plastic-embedded tissue section of a formalin-fixed mouse eye.” This is remarkably close to our best estimates (see below). When removing all aqueous humor from the eye, John et al.^13^ measured a total aqueous volume of 5.8 µL in C57BL/6J mice (n=12, mean ± SEM) and 5.1 ± 0.4 µL in C3HeB/FeJ mice (n = 9), with all mice being 8 to 12 weeks old. Using a similar aspiration method, Zhang et al. ^11^ and Aihara et al.^10^ measured total aqueous volumes of 7-7.2 µL in CD-1 mice (age 4–6 weeks) and of 5.9 µL in NIH white Swiss mice (8-12 weeks of age), respectively.

Attributing approximately half of the total aqueous volume to the anterior chamber, consistent with our data, the above studies would imply anterior chamber volumes of 2.5-3.6 µL, which is larger than our direct optical measurements of 1.55-2.41 µL. Some of this difference may be due to strain and age effects; further, one cannot exclude the possibility of inadvertent collection of secondary aqueous during aspiration, despite careful efforts to avoid such effects^46, 47^. Finally, we note that the Zhang et al. data is larger than the other direct measurements and is perhaps somewhat of an outlier, especially considering that the mice in that study were only 5-6 weeks of age. Thus, we are inclined to consider total aqueous volumes of 3-6 µL as reasonable, depending on age, with corresponding bounds on anterior chamber volume between 1.55 and 2.8 µL, i.e. from our lowest directly measured volume in 2-month-old BALB/cAnNCrl mice to 47% of the presumed upper bound of total aqueous humor volume of 6 µL.

An important consideration in interpretating our data (and others) is that anesthesia affects ocular physiology in several important ways. First, anesthesia is known to affect aqueous inflow rate in both monkeys^48, 49^ and mice^9^ in an anesthesia and time-dependent manner (see below). Second, anesthesia also affects IOP, which in turn affects ocular volume (and thus anterior chamber volume) through an ocular compliance effect. The literature in this area is somewhat contradictory; we here focus only on IOP measurements in mice where awake IOPs measured by TonoLab rebound tonometry were compared to IOPs under ketamine/xylazine anesthesia, since this anesthetic regimen was used during our vis-OCT imaging. Even with this focus, reported IOP changes due to anesthesia are discordant, ranging from a 2.7 mmHg drop at 10 minutes in BALB/cAnNCrl mice^50^ to a 6.4-7.8 mmHg increase in C57BL/6J mice^51^. We were unable to obtain reliable IOP measurements during the vis-OCT imaging process, but we here argue that in any case, anesthesia-induced IOP changes in anterior chamber volume were likely very small, as follows. Sherwood et al. measured mean ocular compliance in control eyes of 11-week-old C57BL/6J mice to be 43-49 nl/mmHg at a reference IOP of 13 mmHg^52^. This means that an IOP change of 5 mmHg would change total ocular volume by 215-245 nl, which is less than 2% of the total volume of the mouse eye. Even in the unlikely scenario that all the volume change of the eye occurred in the anterior chamber, these IOP-associated volume changes would only be of order 10% of our estimated anterior chamber volumes. Thus, this effect is judged to be small and can be safely ignored.

### Implications for aqueous humor dynamics

As noted above, determination of aqueous inflow rate depends on accurate knowledge of anterior chamber volume. Here we reanalyze a recent paper^9^ on this topic in light of our finding that anterior chamber volume is significantly less than total aqueous volume. We specifically consider the work of Toris and colleagues, who used a custom fluorophotometer to estimate an aqueous inflow rate, *Q*, according to a standard equation for human eyes

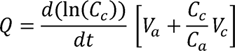

where *C_c_* and *C_a_* are measured concentrations of fluorescein in the cornea and anterior chamber, respectively, and *V_c_* and *V_a_* are the volumes the cornea and anterior chamber, respectively. We have selected this paper not because the measurements were poorly done; quite the converse – the work represents the use of custom technology to carefully determine inflow rates in the mouse eye. We note in passing that there are a number of assumptions underlying the above equation, some of which may be less valid in the mouse eye vs. the human eye, e.g. neglect of tracer diffusion into the posterior chamber. Here we will not concern ourselves with these complex topics, and simply investigate the effects of different anterior chamber volumes.

If we denote the true value of the anterior chamber by 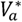, with the corresponding true value of the aqueous outflow rate being denoted by *Q*^*^, then we can write

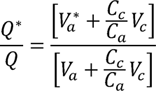

which can be interpreted as a correction factor for reported values of *Q* based on an incorrect anterior chamber volume, *V_a_*.

Toris et al. ^9^ took corneal volume to be *V_c =_* 0.5 *uL* and anterior chamber volume to be *V_a =_* 5.9 *uL*. We digitized Figure 5B of the Toris paper and determined that the mean value of the ratio 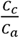 was 4.0. Using this value, we then substituted our range of anterior chamber volumes in the above equation to determine that 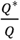 lies in the range 0.45-0.61. Stated differently, the over-estimation in the Toris et al. paper in reported aqueous flow rate is somewhere between 39-55%, which is substantial. Toris et al. reported an aqueous flow rate of 90 ± 70 nl/min in female CD-1 mice greater than 6 months of age and weighing between 35 and 45 g under ketamine/xylazine anesthesia. Using the above correction factor, we would instead estimate a corrected mean aqueous inflow rate of *Q^*^ =* 40 – 55 nl/min. The expected aqueous inflow rate in younger (smaller) mice would be less than the above value. Other studies of inflow rate that make similar assumptions about anterior chamber volume will suffer from the same inaccuracies^3, 8, 10, 11^.

It is important to note that Toris et al. also reported a large effect of anesthesia on aqueous inflow rate in the mouse, with the estimated inflow rate under 2,2,2-tribromoethanol anesthesia being more than 2-fold greater than that estimated under ketamine/xylazine, reinforcing the point that careful consideration of the anesthesia regimen is indicated when studying aqueous humor dynamics ^9^. In what follows, we will consider the case of ketamine/xylazine anesthesia, since this is a commonly used regimen in mice (although different groups use different doses) and because our vis-OCT measurements were obtained on mice under this regimen.

Goldmann’s equation relates inflow rate to other AHD parameters, and may be written as

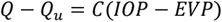

where *EVP* is episcleral venous pressure; *C* is pressure-dependent (conventional) outflow facility; *Q* is aqueous inflow rate; and *Q_u_* is pressure-independent outflow rate, sometimes called unconventional outflow. Sherwood et al. measured facility in 66 *post mortem* eyes of C57BL/6J mice (10- to 14-week-old males), determining a geometric mean population value of 5.89 nL/min/mmHg at an IOP-EVP difference of 8 mmHg, which corresponds to a conventional outflow rate of 47 nL/min. Using this conventional outflow rate with our adjusted inflow rates above indicates that 0-15% of aqueous humor is predicted to exit the eye via the pressure-independent (unconventional) route, similar to that seen in humans and monkeys^53^. Using values for inflow that are not corrected for anterior chamber volume causes this estimate for unconventional outflow to jump to 48%. Regrettably, the above calculations have drawn on data from different strains and ages of mice, and carrying out careful measurements of inflow rate, anterior chamber volume and aqueous outflow facility in mice of the same age, sex and strain may help us better understand the role of unconventional outflow in mice, which has been controversial in the past^53^.

### Limitations

A drawback of our study is that the anterior hyaloid membrane was not well visualized. To address this, we approximated the hyaloid membrane location by the equator of the globe, an assumption we validated with micro-CT imaging. A second limitation is that we could only visualize the posterior chamber in non-pigmented mice, even when using our advanced robotic vis-OCT imaging approach, although we compared estimates from vis-OCT images to standard histology of pigmented mice. Finally, in this study we considered only a few mouse strains and limited ages. Future work should investigate more strains and also a wider range of ages. In fact, chamber volumes could be tracked longitudinally in an *in vivo* setting to assess how specific treatments or procedures impact ocular development and growth.

## Acknowledgments

We thank Prof. Darryl Overby (Imperial College London) for insightful comments on the manuscript and Ying Hao (Duke Eye Center Core Facility) for histological studies. This work was supported in part by National Institutes of Health grants R01EY029121, U01EY033001, R01EY033813, R01EY034740, R01EY034353, R01EY030124, R01EY032062, R01EY032507, F30EY034033, R01EY031710 and R44EY026466, P3EY019007, P30EY005722, Illinois Society for the Prevention of Blindness, the Christina Enroth-Cugell and David Cugell Fellowship for Visual Neuroscience and Biomedical Engineering, unrestricted departmental funding (Columbia) from Research to Prevent Blindness, NY First Empire Fund, and the Georgia Research Alliance (CRE).

## Supplemental Text

### Vis-OCT imaging system

Light from a supercontinuum laser (SuperK EXTREME, NKT Photonics) was filtered using a dichroic mirror (DMSP650, Thorlabs), spectral shaping filter (34-443, Edmund Optics), and bandpass filter (FF02-694/SP-25, Semrock) before being sent to a 90:10 fiber coupler (TW560R2A2, Thorlabs). The reference arm consisted of polarization controllers (FPC560, Thorlabs) and BK7 dispersion compensation glass (27-852, Edmund Optics). Light in the sample arm was scanned using a pair of galvanometer mirrors (Compact-506, ScannerMax) through a 25 mm achromatic doublet scan lens (AC127-025-A, Thorlabs). The light from the scan lens was focused on the sample. Reflected light from the sample arm and the light transmitted through the reference arm was coupled to a second 50:50 fiber coupler (TW560R2F2, Thorlabs). Two spectrometers (Blizzard SR, Opticent Health) operating from 510 nm to 610 nm detected the interferogram signals propagating through the second fiber coupler for image reconstruction. We used two spectrometers for balanced detection to eliminate the influences of relative intensity noise ^54^. The axial resolution of the system is 1.3 µm ^24^, and the lateral resolution is 8.8 µm as measured with a USAF51 target card (R1DS1P, Thorlabs). The vis-OCT’s A-line rate was 75 kHz, and the illumination power on the sample was 0.8 mW.

### Fusion of individual volumes into a composite volume

A total of eight transformations were obtained from the eight vis-OCT sub-volumes. Using these transformations, we mapped the coordinate system of all volumes to the coordinate system of the first acquired vis-OCT volume (Fig. 2d). For vis-OCT volumes not adjacent to the first volume, the transformation matrices for each volume between the given volume and the first volume can be multiplied to determine the net transformation of the given volume to the first volume. Specifically, given the transformation *T_i_* mapping volume *i* onto the reference coordinate system, coordinate (*x_i_, y_i_, z_i_*) in the reference frame of the volume is mapped to (*x’, y’, z’*) in the reference coordinate system by

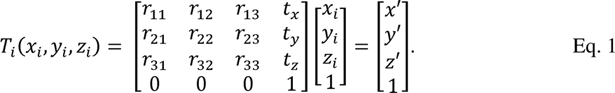

After mapping all eight volumes onto a common reference frame, we identified the overlapping regions between each adjacent volume pair. Next, we applied an iterative closest point (ICP) algorithm to refine the transformation of each volume to the common reference frame^55^. Specifically, we used ICP to minimize the distance between the point clouds of overlapping regions of adjacent volumes. The purpose of the refinement step was to increase the number of points used to register adjacent volumes and to address the loop closure problem ^31^, which results from the error propagating from the multiplication of multiple transformation matrices. After refining the transformation, we defined the intensity of the reconstructed signal in the global reference frame V(*x’,y’,z’*) as

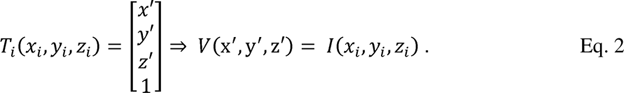

In other words, if *T_i_*maps (*x_i_,y_i_,z_i_*) in the original reference frame of volume *i* to (*x’,y’,z’*), then the intensity of the reconstructed signal V(*x’,y’,z’*) is that of *I*(*x_i_,y_i_,z_i_*) in the original reference frame of volume *i.* For situations in where the mapped pixel was shared between two adjacent scans, i.e. in which *T_i_*maps (*x_i_,y_i_,z_i_*) to (*x’,y’,z’*) and *T_j_* maps (*x_j_,y_j_,z_j_*) to the same (*x’,y’,z’*), we assigned V(*x’,y’,z’*) as max{I(*x_i_,y_i_,z_i_*) in the reference frame of volume I, I(*x_j_,y_j_,z_j_*) in the reference frame of volume j}.

### Determination of the approximate anterior hyaloid membrane location

First, we identified the inner boundaries of the anterior and posterior chambers (Fig. 4a) which coincided with the anterior boundary of the lens (Fig. 4b). Next, we fit an ellipsoid to the lens boundary by minimizing the least-squared error (LSE)

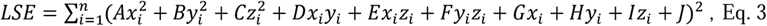

where (x_i_, y_i_, z_i_) are the lens boundary points and A-J are coefficients for the general quadric surface equation (Fig. 4c). After fitting, we found the center of the lens ellipsoid (x, y, z) by solving the following equation.

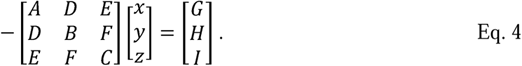

We determined the optical axis of the eye by using principal component analysis ^56^ of the coordinates for all segmented voxels corresponding to the anterior chamber. The calculated principal components are orthogonal vectors, with the vector aligning most closely to the z-axis of the reconstructed volume being the direction of the optical axis of the eye. We approximated the anterior hyaloid membrane as the plane passing through the center of the ellipsoid with a normal vector the same as the optical axis of the eye (Fig. 4d).

Finally, we updated the boundaries of the lens and outer surface of the posterior chamber (Fig. 4e) and applied a k-means-based volumetric segmentation on the full volumetric reconstruction ^57^. We updated the posterior chamber segmentation using the newly segmented posterior chamber outer boundary, the lens boundary, and the plane approximating the anterior hyaloid membrane, as highlighted by the blue regions in Fig. 4f.

## Supplemental Figures

**Supplemental Figure S1:**
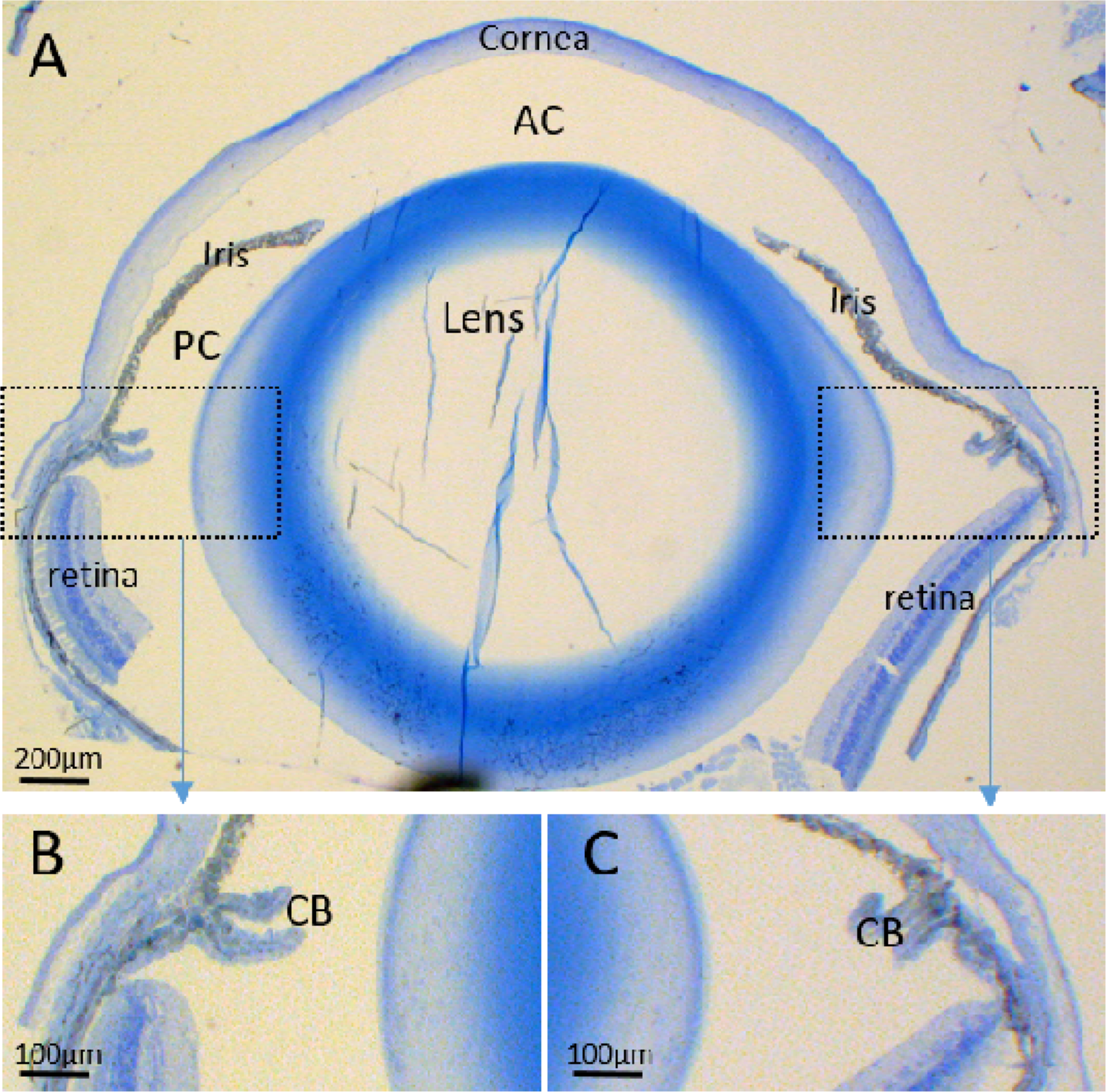
**A)** Histologic cross-section of a mouse eye from a 3-month-old female C57BL/6J wild-type mouse. Although distortion of the globe is evident due to histologic preparation, it is evident that the lens equator coincides approximately with the posterior margin of the ciliary body and the anterior termination of the retina. **B)** and **C)** show zoomed-in views of limbal area showing the left and right sides of the CB and anterior edges of retina. AC: anterior chamber, PC: posterior chamber, CB: ciliary body.

**Supplemental Figure S2:**
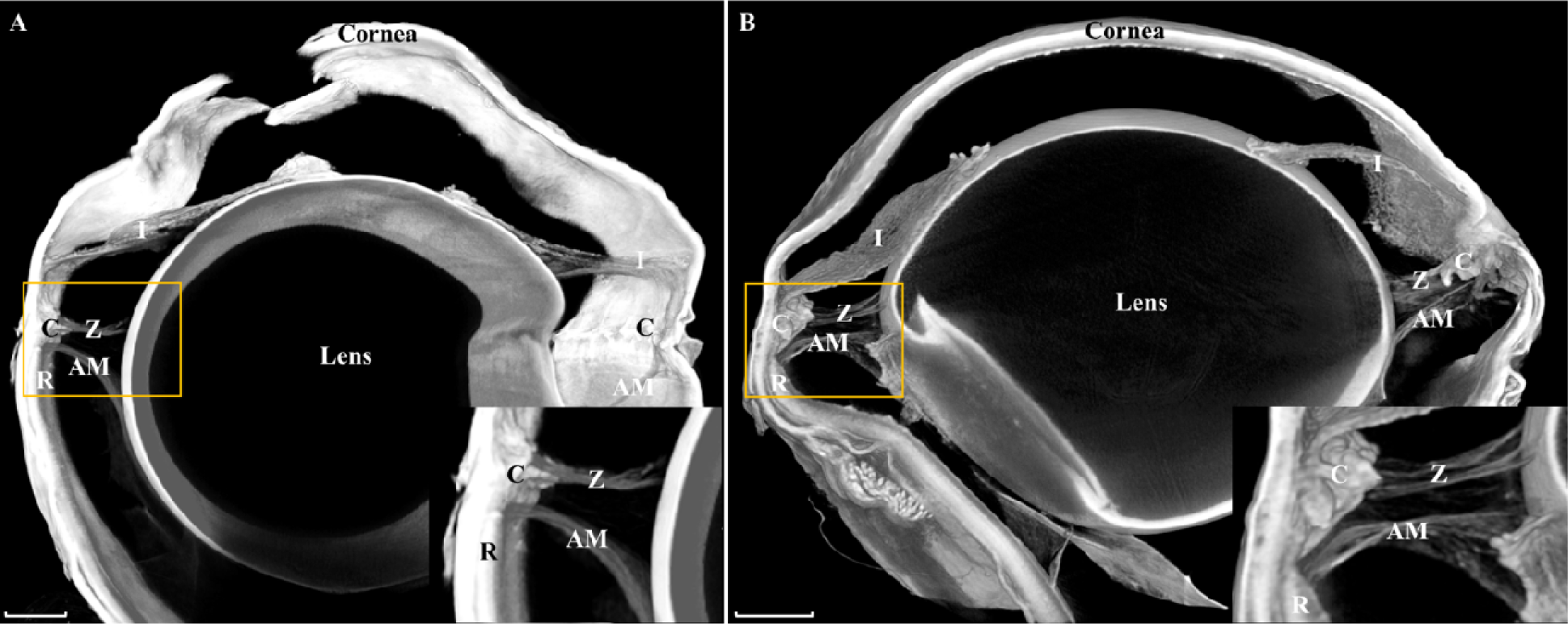
Digital reconstructions from 3-dimensonal micro-CT images of an eye from a 13-month-old DBA/2J female mouse **(A)** and a 1.5-month-old C57BL/6J female mouse **(B)**. Scan resolutions are 0.87 μm and 0.75 μm, respectively. In (A), a corneal window was created to facilitate contrast agent penetration. The far side of the eye has been digitally removed to better visualize the structures of interest. Zonules are seen stretching from the ciliary processes to the lens. The peripheral edge of the anterior hyaloid membrane extends to the anterior boundary of the retina. The yellow boxes outline the areas of the insets. AM = Anterior Hyaloid Membrane, C = Ciliary Body, I = Iris, R = Retina, Z = Zonules, Scale bars = 250 _μ_m.

